# High-Resolution Coastal Blue Carbon Site Intelligence: A Multi-Attribute Geospatial Pipeline for National-Scale Mangrove Assessment

**DOI:** 10.64898/2026.02.20.706974

**Authors:** Jay Gutierrez

## Abstract

The voluntary blue carbon market is severely bottlenecked by outdated methodologies that apply broad, coast-level carbon averages across low-resolution spatial units, systematically failing to account for micro-site ecological realities and critical socio-political constraints. To resolve this structural deficit, this paper introduces the High-Resolution Geographically-Explicit Blue Carbon Assessment (HiGEBCA) pipeline, an innovative geospatial architecture that shifts site intelligence from monolithic raster grids to a topologically verified, hyper-dimensional polygon infrastructure. Operating on 1,601 distinct mangrove features across Colombia, the pipeline mathematically binds 47 ecological attributes to each polygon, integrating Monte Carlo uncertainty propagation, climate-stratified soil organic carbon, and rigorous biodiversity quantification spanning 293 taxon-code pairs. A diagnostic CatBoost machine learning emulator (R² = 0.926) deployed within the pipeline empirically demonstrates that local climate classes and biodiversity metrics drive over 96% of the variance in carbon density, proving that traditional broad biome classifications are inadequate for accurate micro-site valuation. Crucially, the HiGEBCA framework pioneers the integration of operational reality into natural capital assessment. When applied to Colombia’s theoretical national estate of 276,430 hectares (containing an estimated 478 million tCO₂e), the pipeline executes a rigorous REDD+ white space assessment alongside hard mathematical filters for legal land tenure, armed conflict, and regulatory overlap. This strict governance filtration shatters the illusion of massive, easily accessible natural capital, systematically reducing the viable, investment-grade portfolio to a highly de-risked 4,000 to 12,000 hectares. Designed for cross-jurisdictional replication, the HiGEBCA pipeline establishes a new, transparent standard for prioritizing high-integrity blue carbon assets, providing a quantitative mandate for investors seeking to maximize climate impact, capture biodiversity premiums, and definitively mitigate operational risk.

## 1. Introduction

Mangrove ecosystems store three to five times more carbon per hectare than most terrestrial forests (Donato et al. 2011), with 49 to 98% of total ecosystem carbon locked in anoxic soil that persists for centuries when undisturbed (Alongi 2012). They simultaneously contribute substantially to fisheries productivity, buffer storm surges, filter water, stabilize sediment, and support biodiversity across marine, estuarine, and terrestrial taxa.

The global voluntary carbon market has spent two decades attempting to monetize mangrove-derived carbon, yet fewer than 7 million blue carbon credits have been issued globally, representing only a tiny fraction of the voluntary carbon market volume (Farahmand et al. 2025). Moreover, the number of registered blue carbon projects worldwide remains below 100 as of early 2026. The supply constraint is partly methodological. A recent synthesis identified 14 procedural improvements needed to strengthen blue carbon science, noting that differing methodologies introduce cumulative biases causing carbon stock estimates to differ by up to 10-fold across studies (Macreadie et al. 2025). Standard assessments globally rely on remote-sensing-derived extent estimates with coast-level carbon factors. The most comparable published blue carbon site prioritization applied spatial multi-criteria decision analysis (MCDA) with three criteria at the landscape scale without polygon-level biodiversity integration (Rog et al. 2024). To the best of our knowledge, no published assessment combines polygon-level geometry with soil typology, climate stratification, biome-specific carbon parameters, and multi-taxon biodiversity inventories in a single analytical framework.

This paper presents a High-Resolution Geographically-Explicit Blue Carbon Assessment (HiGEBCA) pipeline that addresses these gaps. The HiGEBCA pipeline processes 47 ecological attributes per polygon, propagates carbon estimation uncertainty through Monte Carlo simulation, integrates biodiversity co-benefits from government-source field data, applies Technique for Order of Preference by Similarity to Ideal Solution (TOPSIS) MCDA with nine weighted criteria, assesses REDD+ project overlap to identify white space opportunities for project development, trains a machine learning emulator to identify the ecological drivers of carbon density variation, and designs a field sampling plan optimized for uncertainty reduction. The HiGEBCA pipeline operates on government-source data (Colombia’s 1:100,000-scale National Ecosystem Map) and produces outputs at a resolution that exceeds published blue carbon site assessments: 1,601 individual polygons, each characterized across 47 dimensions, dissolved into 12 coherent Areas of Interest (AOIs) covering 276,430 ha. The contribution of this work is methodological. The HiGEBCA pipeline is designed as replicable infrastructure applicable to any country with equivalent high-resolution ecological mapping, not as a one-off assessment. All analytical outputs, from carbon estimates to priority rankings to field sampling candidates, are reproducible from the source data through documented code. This paper demonstrates the full pipeline using Colombia as the case study, discusses its validation against existing blue carbon projects, and evaluates its scalability to other jurisdictions.

## 2. Data Sources and Study Area

### 2.1 Primary Dataset

The HiGEBCA pipeline operates on one of Colombia’s most authoritative ecological dataset: the IDEAM/SINCHI/IAvH/INVEMAR 1:100,000-scale National Ecosystem Map, 2024 edition (IDEAM et al. 2024). This is the official government source referenced during carbon project validation by several registries, which is not a mere satellite-derived approximation or modeled estimate. From this dataset, 1,601 individual mangrove features were extracted using the filter u_sintesis contains ‘manglar’ (case-insensitive), reprojected from EPSG:4686 (MAGNA-SIRGAS geographic, the coordinate reference system, CRS) to EPSG:3116 (MAGNA-SIRGAS Colombia Bogota zone) for area calculations, and dissolved by hydrographic zone (nom_zh attribute) to produce 12 coherent AOIs covering 276,430 ha.

### 2.2 The 47-Attribute Schema

Each of the 1,601 features carries 47 ecological attributes, of which standard blue carbon assessments use five or fewer. The attributes span seven analytical dimensions:

This attribute depth enables polygon-level ecological characterization that distinguishes, for example, an estuarine deltaic mangrove stand on hydromorphic soils in a superhumid climate (high soil carbon accumulation) from a fringe mangrove on sandy substrate in an arid climate (lower soil carbon), even when both fall within the same hydrographic zone. Arguably, standard assessments applying a single coast-level carbon factor cannot make this distinction.

### 2.3 Complementary Datasets

The HiGEBCA pipeline integrates two additional geospatial datasets for the REDD+ white space assessment. First, the Verra Verified Carbon Standard (VCS) Registry provides project boundaries for registered REDD+ and blue carbon projects in Colombia; one mangrove project was identified (Vida Manglar, VCS 2290, Cispata Bay). Second, the Colombian National Registry of Protected Areas (RUNAP) contributes 44 coastal protected area boundaries totaling 8.3 million ha.

### 2.4 Study Area

Colombia’s mangroves span two coastlines: the Pacific (219,839 ha, 79.5%) and the Caribbean (56,591 ha, 20.5%), distributed across 12 hydrographic zones ranging from 137 ha (Islas Caribe) to 70,961 ha (Patia). The Pacific coast is dominated by large estuarine deltaic systems with high soil carbon accumulation; the Caribbean coast features smaller, more fragmented patches with lower carbon density but higher accessibility. This structural asymmetry between ecological value and operational accessibility is a central finding of the HiGEBCA pipeline analysis.

## 3. Methodology

### 3.1 HiGEBCA Processing Pipeline

Raw polygon features were extracted from the national ecosystem map, filtered for mangrove ecosystems, and reprojected to EPSG:3116 for metric area calculations. Features were dissolved by hydrographic zone to produce 12 AOIs. Each AOI inherits aggregated statistics (area-weighted means, polygon counts, diversity indices) from its constituent polygons. All geometric operations used the GeoPandas library with EPSG:3116 projection, which introduces less than 1% area distortion across mainland Colombia (up to 1.7% for offshore islands).

### 3.2 Carbon Stock Estimation

Carbon stocks were estimated using a three-pool Tier 1 model, which was propagated through Monte Carlo simulation (N = 10,000 iterations, seed = 42). Above-ground biomass (AGB) densities were assigned based on bioma_IAvH classification, using published values from Colombian and pan-tropical literature (range: 85 to 220 Mg/ha depending on biome; Komiyama et al. 2008). Fourteen distinct biome classifications produce 14 AGB lookup values. Distributions are lognormal (right-skewed, zero-bounded), reflecting field measurement distributions. Below-ground biomass (BGB). Derived from AGB using published root-to-shoot ratios: 0.39 for Pacific coast, 0.49 for Caribbean coast, reflecting the distinct allometric relationships documented for mangroves in these hydrological regimes. Soil organic carbon (SOC), estimated to a 1-meter depth in accordance with the IPCC 2013 Wetlands Supplement convention, utilizing climate-stratified values derived across six climatic classes from the ‘clima’ attribute.

This stratification captures the moisture gradient that drives soil carbon accumulation: arid Caribbean zones store less soil carbon than superhumid Pacific estuaries, consistent with global controls on mangrove soil carbon identified by Rovai et al. (2018) and Sanderman et al. (2018). SOC values are consistent with Neotropical reference ranges of 250 to 450 Mg C/ha for Colombian mangroves. Finally, a carbon fraction of 0.47 was applied (IPCC Tier 1 default). Each Monte Carlo iteration samples independently from the distribution of each pool, producing per-polygon carbon density estimates with full uncertainty quantification. Zone-level statistics are computed by aggregating polygon-level results.

### 3.3 Landscape Ecology Metrics

The HiGEBCA pipeline computes landscape metrics analogous to those produced by FRAGSTATS for each Area of Interest (AOI). Core Area Integrity (CAI) quantifies the proportion of mangrove area classified as interior forest, defined as being buffered 50 m from any edge; a high CAI value indicates structurally intact and unfragmented stands. The fragmentation risk index is a composite metric derived from patch density, edge density, area coefficient of variation, and the mean weighted shape index. The integral index of connectivity (IIC) is a graph-theoretic metric computed at four distance thresholds (500 m, 1,000 m, 2,000 m, 5,000 m), quantifying the degree to which patches function as a connected network rather than isolated fragments. The isolation index, based on nearest-neighbor distances and stepping stone availability, identifies patches that cannot exchange propagules or fauna with neighboring mangrove stands.

### 3.4 Biodiversity Assessment

Biodiversity quantification operates at three levels: alpha diversity (within-zone richness), beta diversity (between-zone turnover), and portfolio-level complementarity. Alpha diversity was quantified by extracting species richness counts for five taxonomic groups (amphibians, birds, angiosperms, mammals, and reptiles) from the coded attributes of each polygon. These attributes are structured as hyphen-separated strings representing species assemblages. Area-weighted richness counts were subsequently calculated for each zone and normalized to generate a composite biodiversity index ranging from 0.0 to 1.0.

Beta diversity was assessed by computing pairwise Jaccard dissimilarity matrices across all 12 zones, utilizing 293 taxon-code pairs. This dissimilarity was subsequently partitioned into turnover and nestedness components, following the methodology proposed by Baselga (2010). A high degree of turnover between zones suggests the presence of distinct species assemblages, indicating a high conservation irreplaceability for each zone. Conversely, a high nestedness component signifies that species-poor assemblages constitute proper subsets of species-rich assemblages.

Species complementarity is assessed through an exhaustive evaluation of pairwise and triplet zone combinations to determine cumulative species capture. This analysis identifies the minimal configuration of zones necessary to maximize coverage of the national species pool, thereby providing direct guidance for portfolio design intended to secure biodiversity co-benefit certification, such as that offered under the Climate, Community & Biodiversity Standards (CCB) or the Sustainable Development Verified Impact Standard (SD VISta).

The Carbon-Biodiversity Pareto frontier is delineated by plotting zones within the multidimensional space defined by carbon density and a biodiversity index. The Pareto-optimal front represents a set of zones that achieve the maximum possible performance across both objectives simultaneously. Specifically, any zone on this frontier is characterized by the property that no other zone exists that is superior in both carbon density and biodiversity index; thus, it is undominated in both dimensions.

### 3.5 Multi-Criteria Decision Analysis

The HiGEBCA pipeline applies TOPSIS with nine weighted criteria:

Weights sum to 1.00. Criteria values are normalized using vector normalization before computing Euclidean distances to the positive and negative ideal solutions. TOPSIS scores are bounded [0, 1], with higher scores indicating greater proximity to the ideal solution across all weighted criteria simultaneously. Weight selection follows expert judgment informed by the relative importance of carbon (primary market driver), area (viability threshold), and ecological condition metrics (co-benefit potential). The assigned weights represent an initial, heuristically-derived configuration based on the relative conceptual importance of each factor to site potential, informed by current ecological understanding. This weighting scheme is currently hypothetical and serves as a starting point only, but it is not empirically optimized. We recognize that empirical optimization, ideally through calibration against ground-truthed site success metrics, is a necessary future step to refine the model’s predictive accuracy.

**3.6 REDD+ White Space Assessment**

The white space assessment identifies mangrove areas without existing REDD+ project claims or regulatory exclusions. The analysis overlays the 1,601-polygon mangrove baseline against two constraint layers: (1) Verra VCS project boundaries for registered REDD+ and blue carbon projects in Colombia, where one mangrove project was identified (Vida Manglar, VCS 2290, a grouped Conservation International/South Pole project in Cispata Bay); and (2) RUNAP protected areas (44 coastal sites, protected area polygons classified as medium additionality pending governance verification). Each polygon is classified using a priority hierarchy: CONTESTED (direct REDD+ overlap) > ADJACENT_LEAKAGE (within 5 km buffer) > PROTECTED_MEDIUM_ADDITIONALITY > WHITE_SPACE. Zone-level statistics are aggregated from polygon classifications. The assessment correctly identified 11 mangrove polygons (3,132 ha) in the Sinu zone as contested, consistent with Vida Manglar’s documented project extent in the Gulf of Morrosquillo.

### 3.7 Machine Learning Baseline Emulator

A CatBoost gradient boosting model was developed to predict polygon-level carbon density. The model was trained using 12 ecological features, comprising six continuous and six categorical variables. CatBoost’s native handling of categorical features eliminated the need for one-hot encoding. The continuous features included: area in hectares (area_ha), and counts of various faunal and floral groups (no_anfibio, no_aves, no_magnoli, no_mamifer, no_reptile). The categorical features used were: bioma_IAvH, clima, paisaje, relieve, suelos, and u_sintesis.

Cross-validation employed GroupKFold with $k=5$ groups, delineated by hydrographic zone, to mitigate data leakage arising from spatial correlation among polygons. Four candidate models were evaluated: CatBoost, XGBoost, Random Forest, and Ridge regression. To quantify feature importance, SHapley Additive exPlanations (SHAP) TreeExplainer was applied to the fully trained CatBoost model, reporting feature importance as the mean absolute SHAP value across all polygons. Furthermore, CatBoost quantile regression models were trained at the 10th and 90th percentiles to establish per-polygon uncertainty bounds (P10/P90) on carbon density predictions.

Importantly, the emulator’s purpose is diagnostic rather than predictive: it identifies which ecological attributes most strongly drive carbon density variation across the polygon population, informing both field sampling prioritization and the design of future remote sensing pipelines.

### 3.8 Field Sampling Design

A stratified maximin sampling algorithm was used to select 50 candidate polygons for field verification from the 1,601-polygon population. Selection is priority-weighted across three criteria: emulator uncertainty (weight 0.4), ecological rarity (weight 0.3), and biodiversity richness (weight 0.3). The algorithm enforces proportional representation across all 12 hydrographic zones and the ecological clusters (Figure 2), which were identified through an unsupervised analysis involving Uniform Manifold Approximation and Projection (UMAP) dimensionality reduction followed by Hierarchical Density-Based Spatial Clustering of Applications with Noise (HDBSCAN).

**Figure 1.**
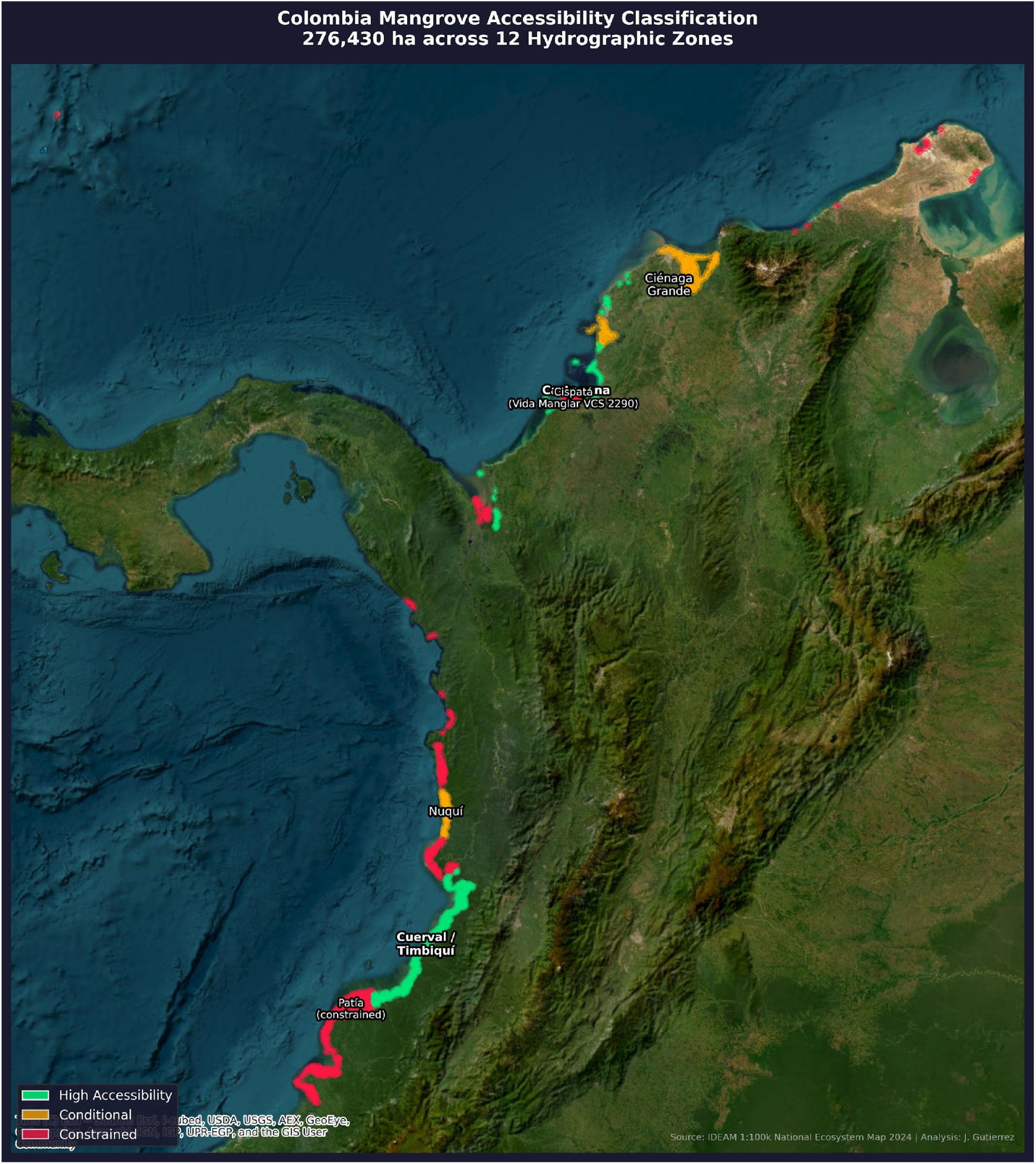
National Mangrove Distribution and Accessibility Classification. Satellite basemap with mangrove extent colored by accessibility classification across 12 hydrographic zones. Green: High Accessibility sites with established community governance and stable security. Amber: Conditional sites where external factors constrain entry. Red: Constrained sites with active REDD+ claims, armed conflict, or undefined carbon rights. Colombia’s 276,430 ha of mangrove span the Caribbean coast (north, 56,591 ha, 20.5%) and Pacific coast (west, 219,839 ha, 79.5%). Source: IDEAM 1:100,000-scale National Ecosystem Map 2024, Verra VCS registry, RUNAP protected areas. CRS: EPSG:4686.

**Figure 2.**
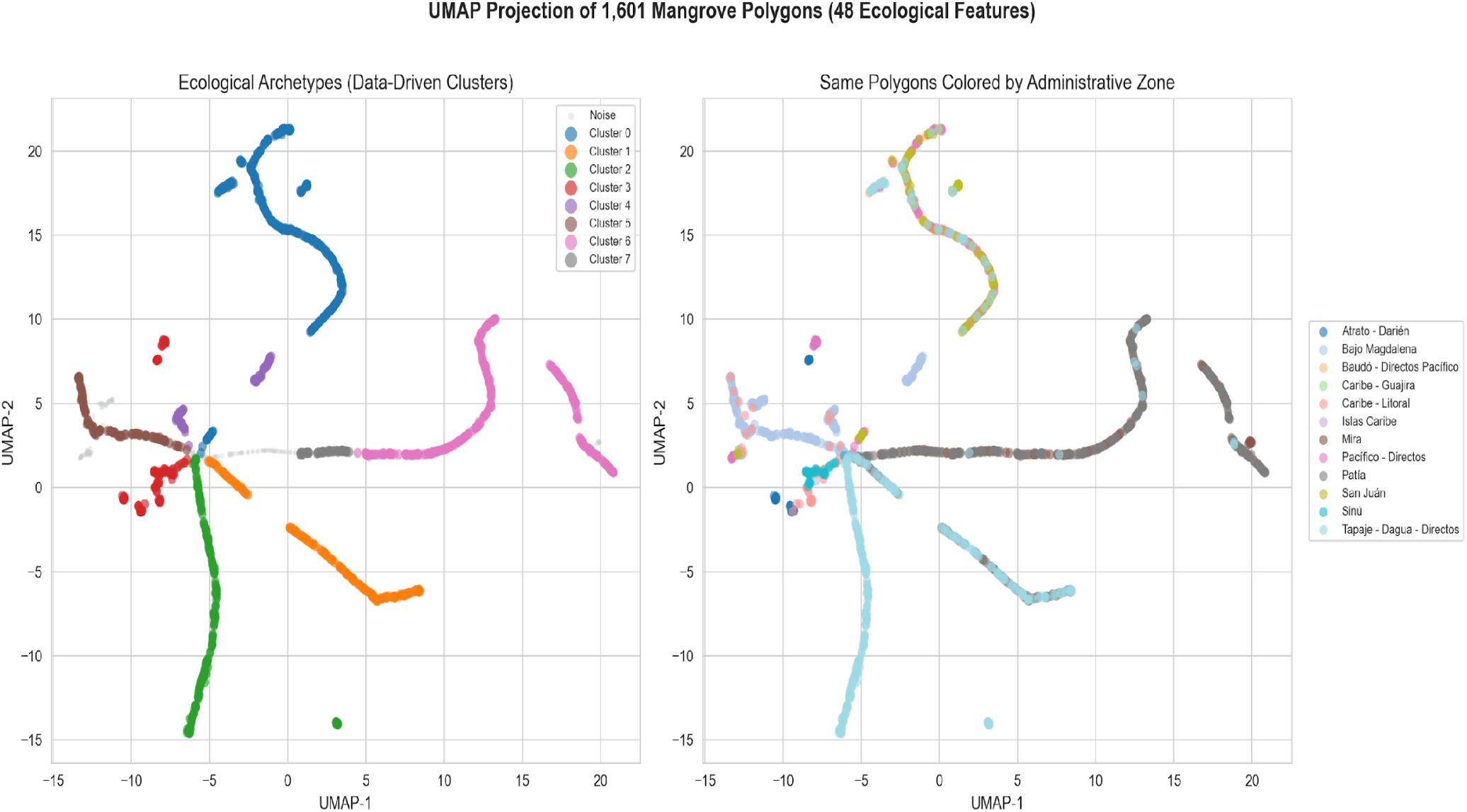
UMAP Ecological Clustering of 1,601 Mangrove Polygons. Scatter plot in two-dimensional UMAP embedding space (n_neighbors = 30, min_dist = 0.3) with points colored by HDBSCAN cluster assignment (min_cluster_size = 40). Feature space: 48 dimensions (6 continuous ecological metrics + 42 one-hot encoded categorical variables from 4 attribute groups). The analysis identifies 8 ecological clusters plus 164 noise polygons (10.2%). Cluster separation tracks the Pacific-Caribbean divide and the moisture gradient.

### 3.9 Governance Constraint Integration

The HiGEBCA pipeline incorporates governance, security, and regulatory constraints as integral, explicit analytical layers, rather than treating them as post-hoc qualifications. This methodological approach establishes a critical distinction between theoretical ecological potential (derived from empirical data) and operational accessibility (the feasibility of implementation with integrity). In the Colombian context, the legal framework significantly influences site accessibility through several mechanisms. Specifically, Constitutional Court ruling T-248 (25 June 2024) mandated that carbon projects creating intense social, cultural, or environmental impacts must adhere to the highest standard of free, prior, and informed consent (FPIC). This mandate was further strengthened in May 2025, when the government instituted a cultural objection right, permitting the revocation of consent. Furthermore, the allocation of carbon rights in mangroves remains legally ambiguous: while Colombian law links carbon rights to land ownership, mangroves are simultaneously classified as public goods where private tenure is strictly prohibited.

The HiGEBCA pipeline classifies sites into accessibility tiers based on governance readiness, security conditions, and regulatory overlap. This classification is itself a methodological output: it demonstrates how constraint filtering can reduce a nominal 276,430 ha portfolio to a subset of 4,000 to 12,000 ha that meets integrity standards, and quantifies what is excluded and why.

## 4. Results

### 4.1 National Mangrove Characterization

The HiGEBCA pipeline has characterized 276,430 hectares (ha) of mangrove ecosystems distributed across 12 distinct hydrographic zones (Figure 3). The spatial distribution of this coverage is highly concentrated: the Patia zone alone encompasses 70,961 ha, representing 25.7% of the total national mangrove area. Furthermore, the four largest zones on the Pacific coast (Patia, Tapaje-Dagua-Directos, Mira, and Pacifico-Directos) collectively account for 180,993 ha (65.5%). In total, Pacific coast mangroves constitute the majority, comprising 219,839 ha (79.5%), while Caribbean coast mangroves total 56,591 ha (20.5%).

**Figure 3.**
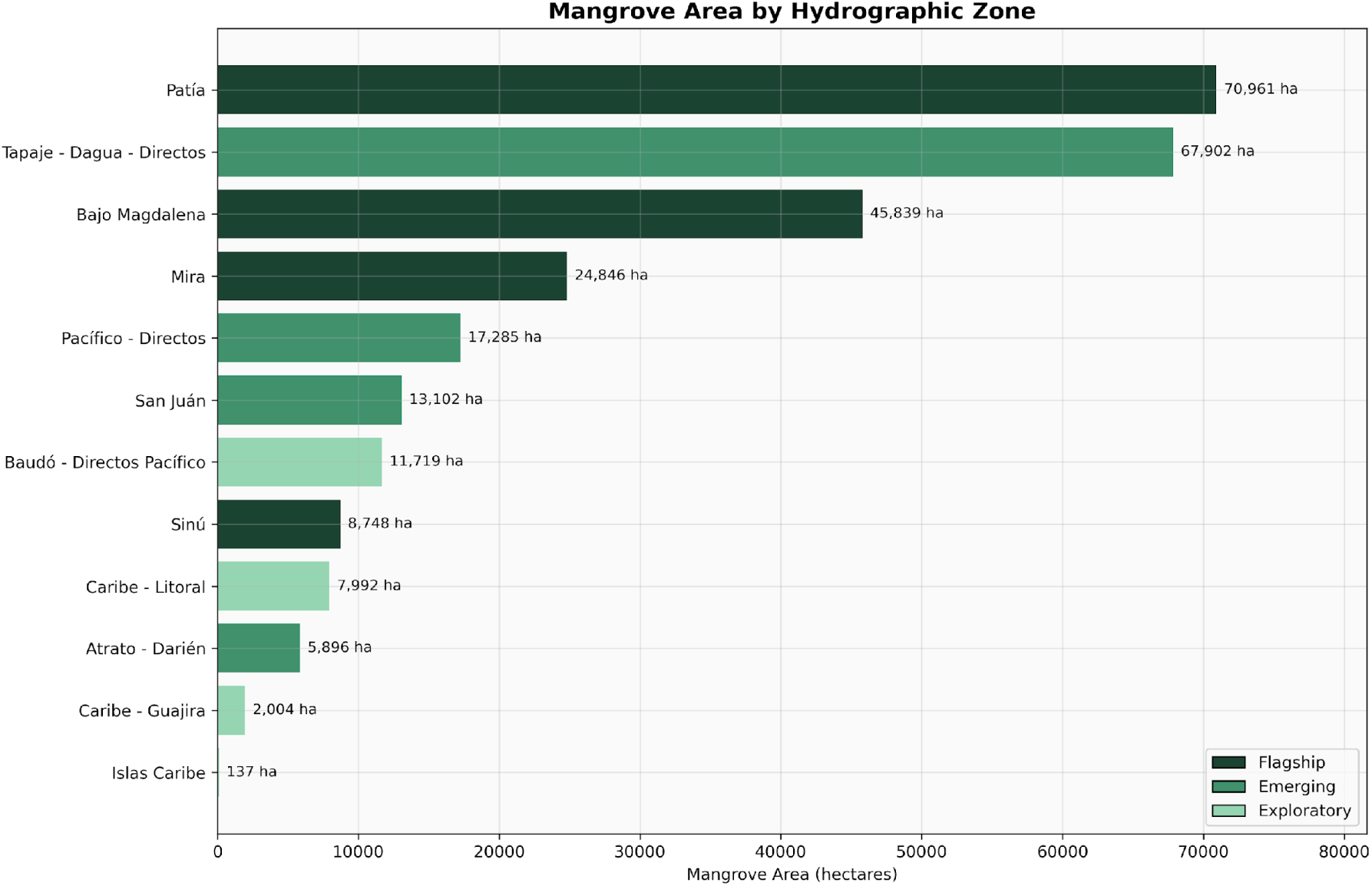
Mangrove Area by Hydrographic Zone. Bar chart showing the area distribution across 12 zones. Patia (70,961 ha, 25.7%) and Tapaje-Dagua-Directos (67,901 ha, 24.6%) together account for 50.2% of the national total. Color coding distinguishes Pacific and Caribbean zones. Source: IDEAM 1:100,000-scale National Ecosystem Map 2024, area calculated in EPSG:3116.

Core area integrity ranges from 56.9% (Islas Caribe, a small island system dominated by edge habitat) to 77.4% (Patia, a large contiguous estuarine complex). The national mean is 68.4%. Fragmentation risk is lowest in the large Pacific systems (Patia: 0.12, Pacifico-Directos: 0.21) and highest in the small Caribbean systems (Caribe-Guajira: 0.52, Bajo Magdalena: 0.64).

IIC at 2 km threshold varies from 0.091 (Caribe-Litoral, highly fragmented urban-adjacent patches) to 0.786 (Islas Caribe, a two-patch system with high internal connectivity but no external connectivity). Among larger systems, Sinu shows the highest connectivity (0.497), reflecting its concentrated patch configuration around the Gulf of Morrosquillo.

### 4.2 Carbon Stock Estimates with Uncertainty

Total ecosystem carbon density ranges from 255.4 Mg C/ha (Caribe-Guajira, an arid Caribbean zone) to 528.1 Mg C/ha (Tapaje-Dagua-Directos, a superhumid Pacific estuarine system) (Figure 4). The national total is 478 million tCO₂e (95% CI: 442M to 520M tCO₂e).

**Figure 4.**
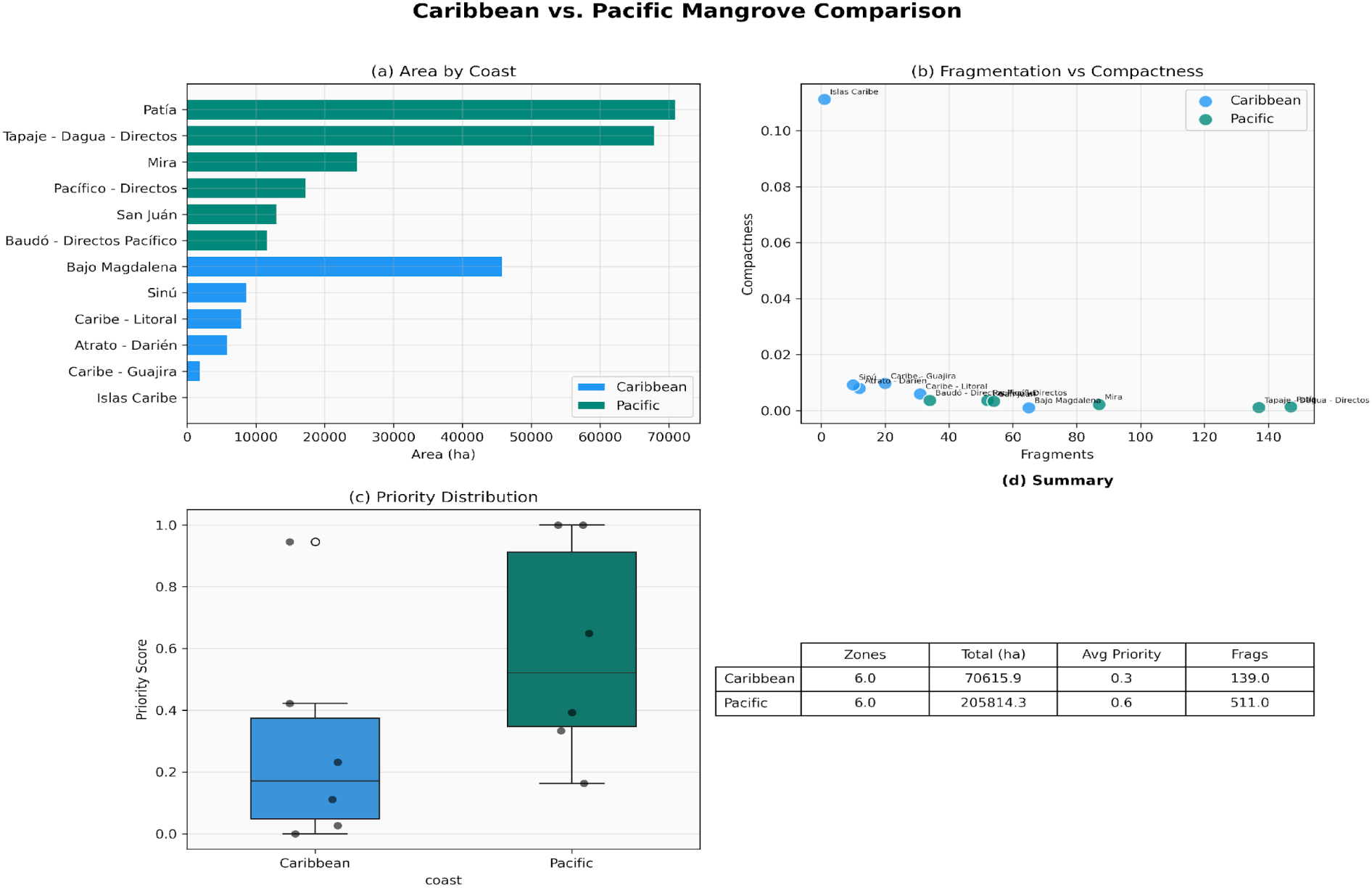
Caribbean vs. Pacific Coast Comparison. Grouped comparison chart showing the structural asymmetry between Colombia’s two coastlines across multiple analytical dimensions. The Caribbean offers smaller, more accessible patches with lower carbon density (255.4 to 441.7 Mg C/ha). The Pacific holds larger systems with higher ecological integrity and higher carbon density (497.6 to 528.1 Mg C/ha).

**Figure 5.**
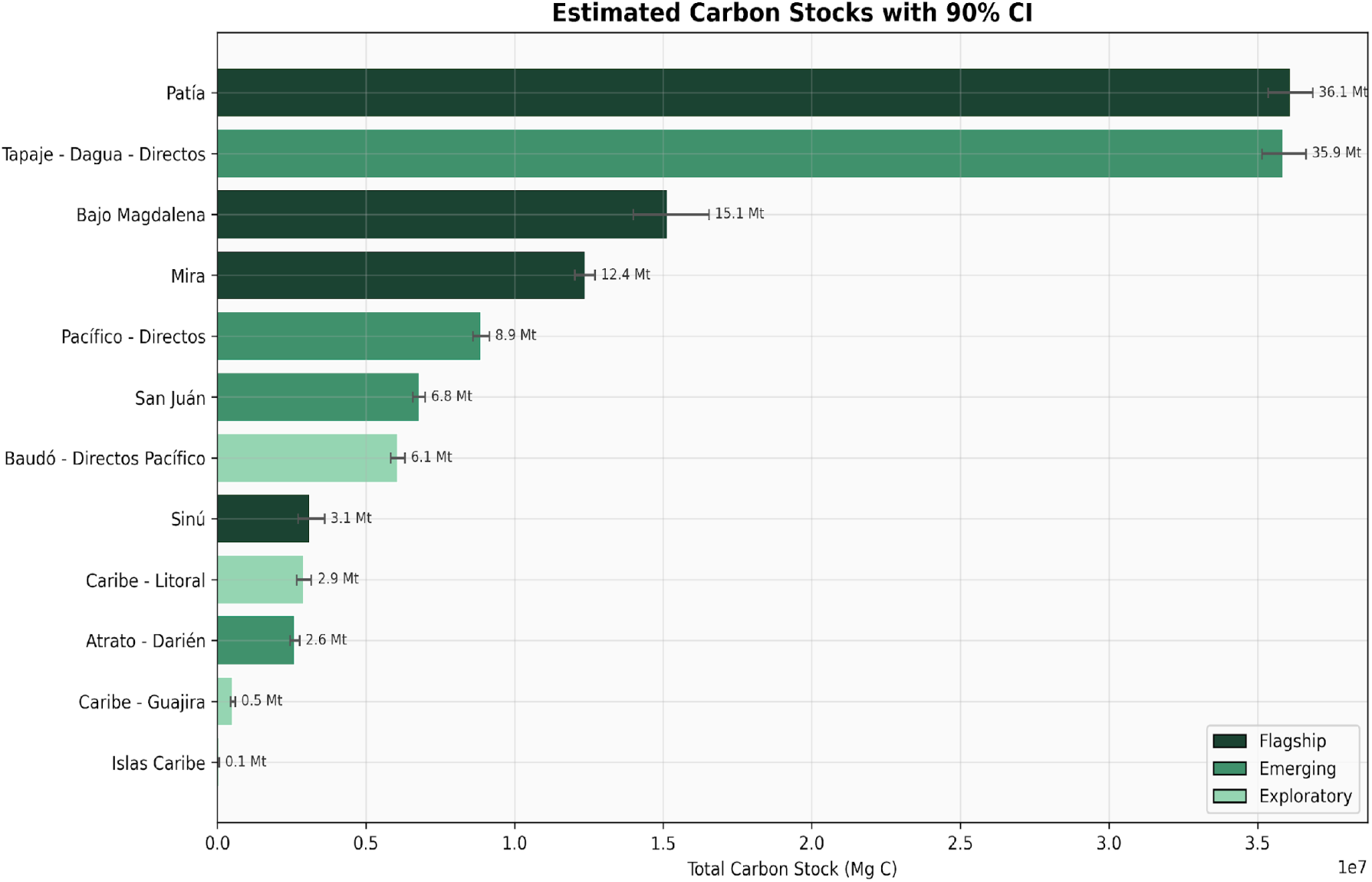
Monte Carlo Carbon Stock Estimates with 95% Confidence Intervals. Bar chart with error bars showing carbon density by hydrographic zone. Monte Carlo simulation (N = 10,000 iterations) propagates uncertainty through three carbon pools: AGB (biome-specific across 14 lookup values from Komiyama et al. 2008), BGB (coast-differentiated root-to-shoot ratios of 0.39 Pacific / 0.49 Caribbean), and SOC (climate-stratified across 6 classes, 200 to 420 Mg C/ha). Error bars: 95% CI. National total: 478 million tCO₂e (95% CI: 442M to 520M).

Interestingly, the climate stratification drives a clear Pacific-Caribbean gradient. All six Pacific zones exceed 497 Mg C/ha; all six Caribbean zones fall below 442 Mg C/ha. SOC accounts for the majority of this difference: superhumid Pacific estuaries accumulate up to 420 Mg C/ha to 1 m depth, compared with 200 Mg C/ha in arid Caribbean zones. This pattern is consistent with the global moisture-SOC relationship documented by Rovai et al. (2018).

### 4.3 Biodiversity Patterns

Biodiversity follows a latitudinal and moisture gradient. The two highest-scoring zones (Mira and Patia, both biodiversity index 1.0, see Figure 6) are Pacific estuarine systems with the highest species richness across all five taxonomic groups: 21 amphibian, 28 bird, 29 angiosperm, 27 mammal, and 27 reptile taxon-codes per area-weighted polygon. The lowest-scoring mainland zone (Caribe-Guajira, 0.22) reflects the arid Caribbean environment’s reduced species carrying capacity. Beta diversity analysis reveals high turnover between Caribbean and Pacific assemblages. Jaccard dissimilarity between Caribbean and Pacific zone pairs ranges from 0.66 to 0.99, indicating that the two coastlines harbor largely distinct species pools (see Figure 7). Within each coast, dissimilarity is lower (0.23 to 0.45 for Caribbean pairs; 0.25 to 0.68 for Pacific pairs), indicating greater compositional similarity among zones sharing a coastline.

**Figure 6.**
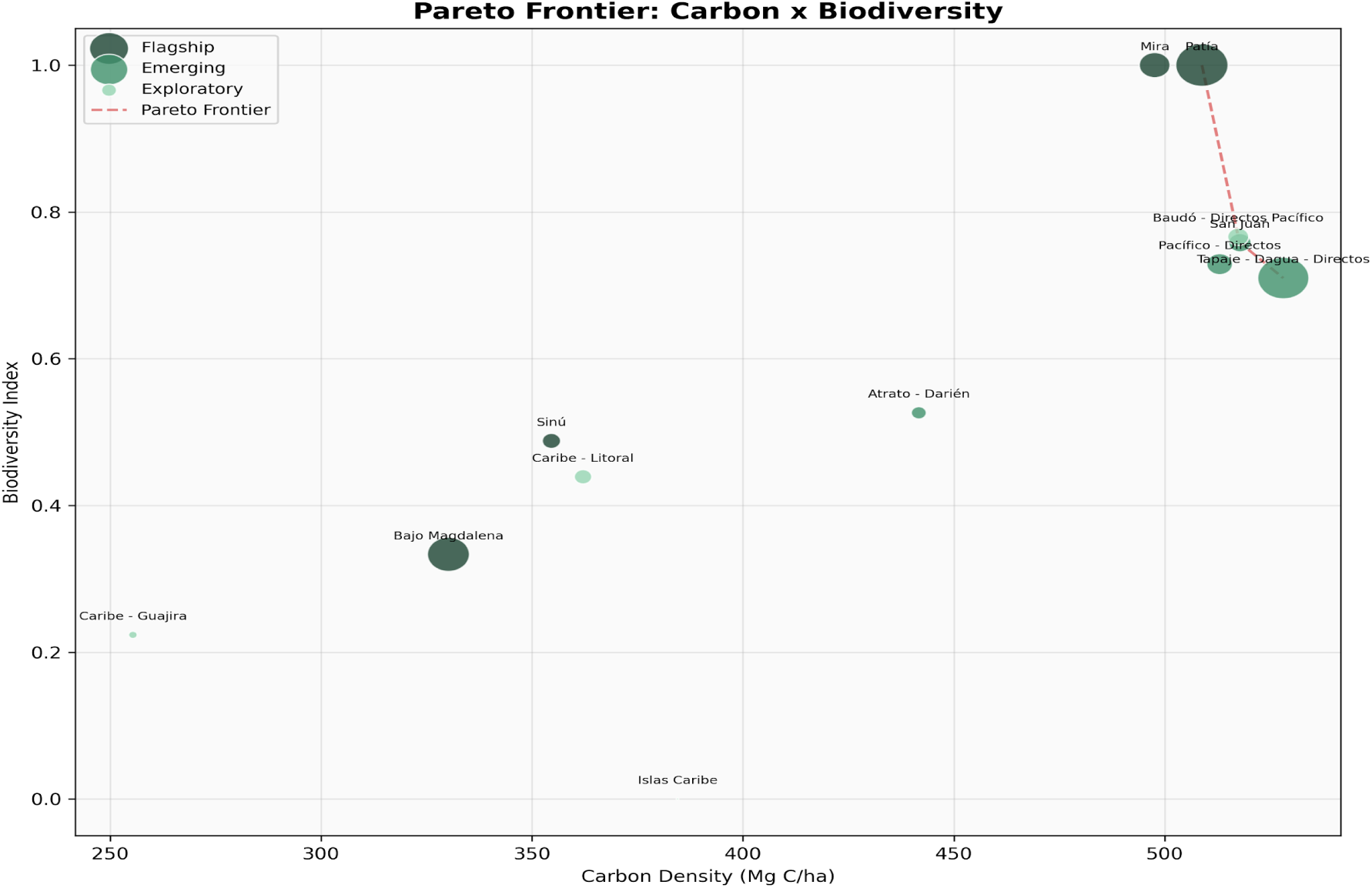
Carbon-Biodiversity Pareto Frontier. Scatter plot with carbon density (Mg C/ha, x-axis) vs. composite biodiversity index (0 to 1, y-axis) for all 12 hydrographic zones. Pareto-optimal zones (upper-right envelope): Patia, San Juan, Tapaje-Dagua-Directos, Baudo-Directos. The two highest biodiversity zones (Mira and Patia, both index = 1.0) share identical species pools (Jaccard dissimilarity = 0.0).

**Figure 7.**
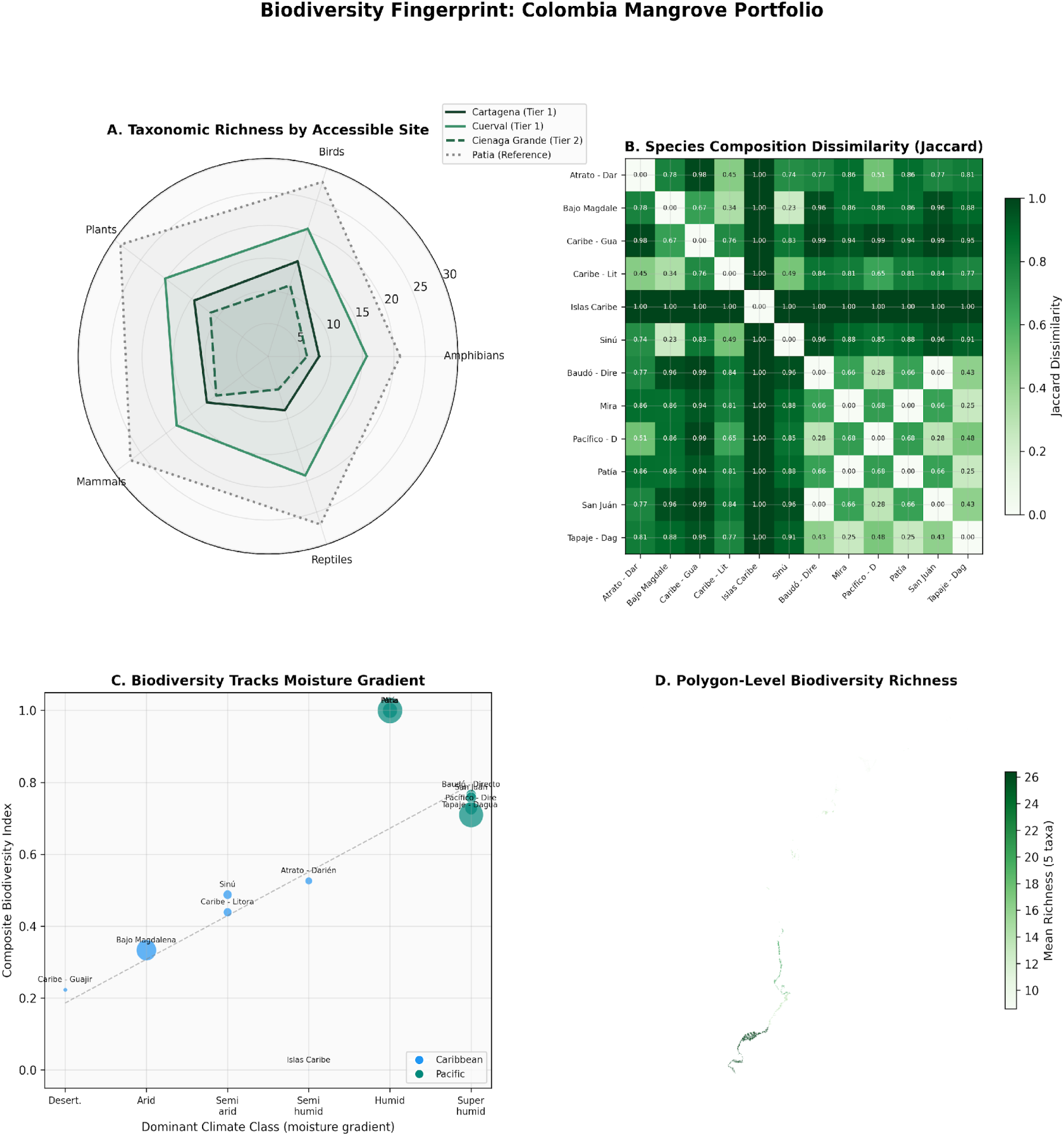
Multi-Panel Biodiversity Fingerprint. (A) Taxonomic richness profiles for each zone across 5 groups. (B) Species composition dissimilarity (Jaccard index) between all 12 zones. (C) Biodiversity richness tracks the moisture gradient. (D) Polygon-level biodiversity richness revealing sub-AOI heterogeneity. Source: IDEAM 1:100,000-scale National Ecosystem Map 2024.

Species complementarity analysis (Figure 7) demonstrates that two zones (Caribe-Litoral and Tapaje-Dagua-Directos) together capture 85.3% (250 of 293 taxon-code pairs) of the national mangrove species pool. This is the optimal two-zone combination. The maximum single-zone coverage is 60.1% (Tapaje-Dagua-Directos alone, 176 taxon-code pairs). A two-zone portfolio spanning both coastlines captures more biodiversity than any set of zones confined to a single coast.

### 4.4 TOPSIS Priority Ranking

The TOPSIS analysis produces a composite ranking across all nine criteria (Figure 8):

**Figure 8.**
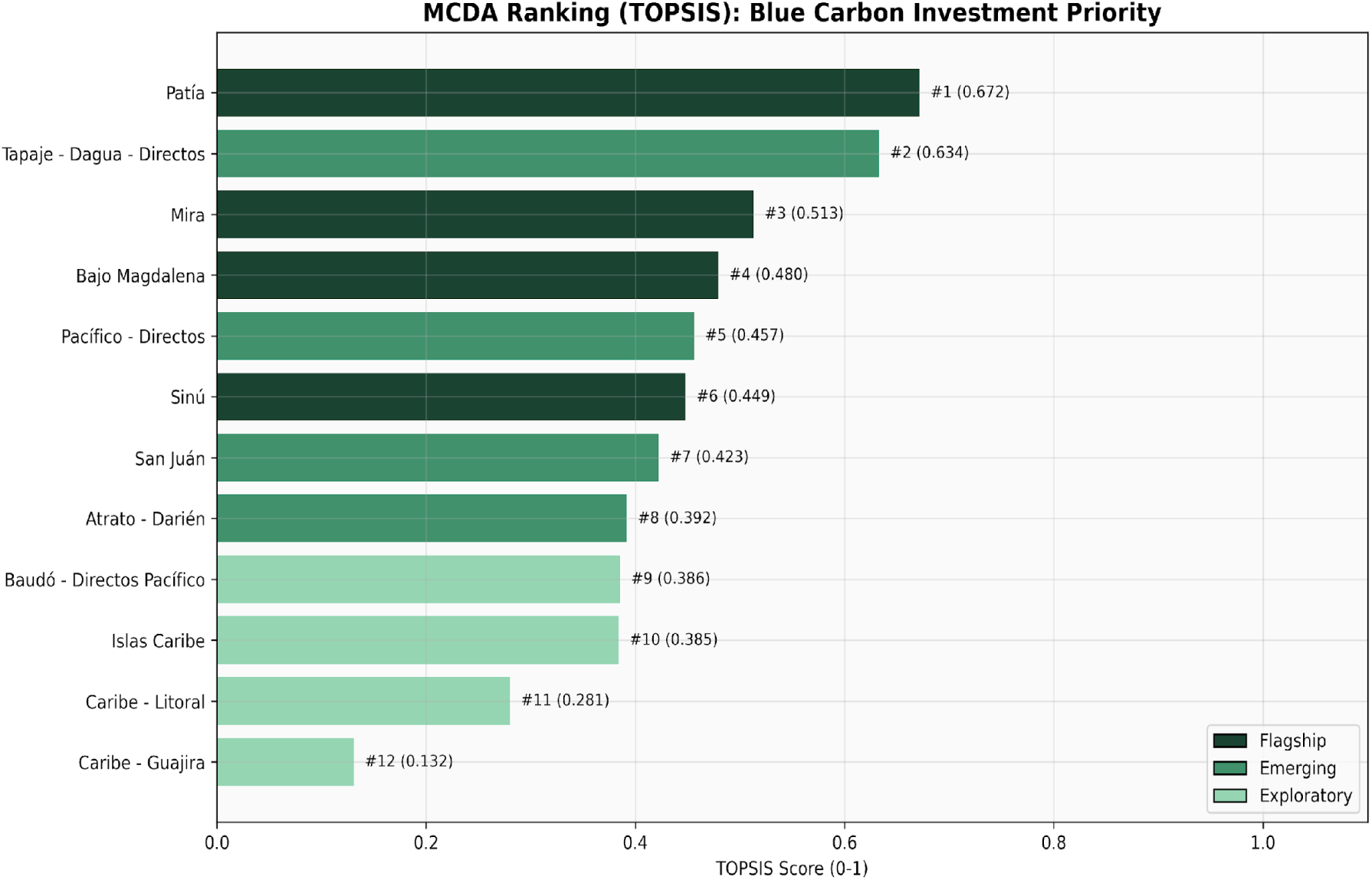
TOPSIS Multi-Criteria Priority Ranking. Horizontal bar chart showing TOPSIS scores (0 to 1) for 12 zones. Nine weighted criteria (Table 3). Patia scores highest (0.672). Scores reflect ecological and carbon potential only; they do not incorporate governance or security constraints.

**Table 1.** Seven analytical dimensions of the 47-attribute schema.

| Dimension | Attributes | Cardinality |
| --- | --- | --- |
| Ecosystem synthesis | `u_sintesis` | 75 types |
| Soil typology | `suelos` | 37 categories |
| Biome classification (IAvH) | `bioma_IAvH` | 14 halobiomes/helobiomes |
| Climate stratification | clima | 6 classes |
| Landscape geomorphology | paisaje | 8 types |
| Relief characterization | relieve | 14 categories |
| Biodiversity (5 taxonomic groups) | anfibios, aves, flores, mamíferos, reptiles | 293 taxon-code pairs |

**Table 2.**
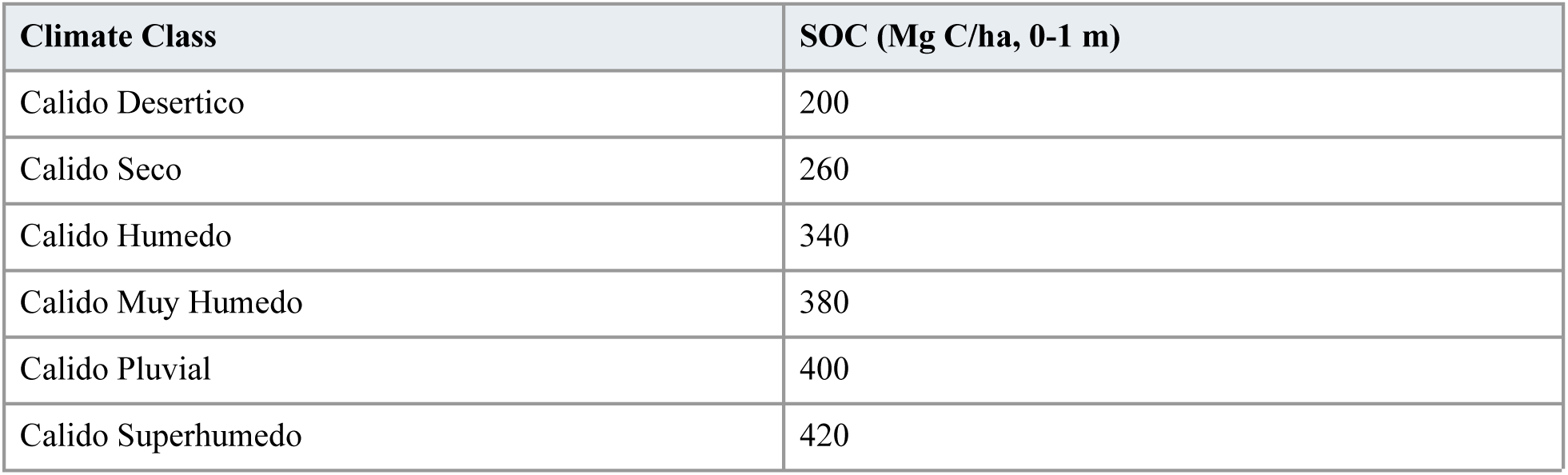
Climate-stratified soil organic carbon values.

**Table 3.** TOPSIS criteria, weights, and optimization direction.

| Criterion | Weight | Direction | Rationale |
| --- | --- | --- | --- |
| Total carbon stock | 0.25 | Maximize | Primary climate value |
| Total area | 0.15 | Maximize | Scale for project viability |
| Core area integrity | 0.10 | Maximize | Ecological condition indicator |
| Biodiversity index | 0.10 | Maximize | Co-benefit certification potential |
| Integral connectivity (2 km) | 0.10 | Maximize | Ecosystem resilience |
| Fragmentation risk | 0.10 | Minimize | Degradation vulnerability |
| Flagship status | 0.10 | Maximize | Existing project/literature validation |
| Isolation risk | 0.05 | Minimize | Connectivity to broader landscape |
| Biome diversity (Shannon) | 0.05 | Maximize | Ecological heterogeneity |

**Table 4.**
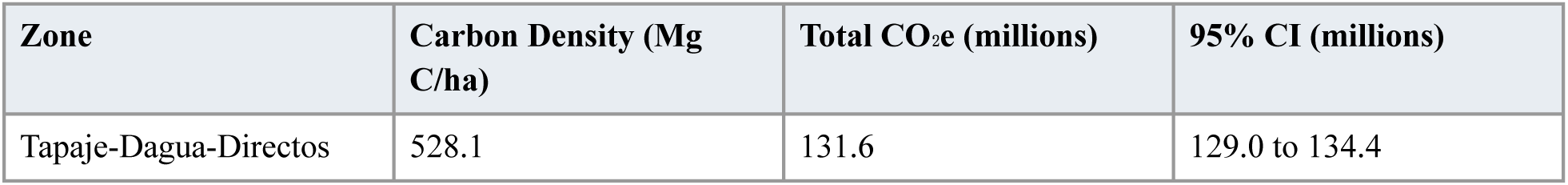
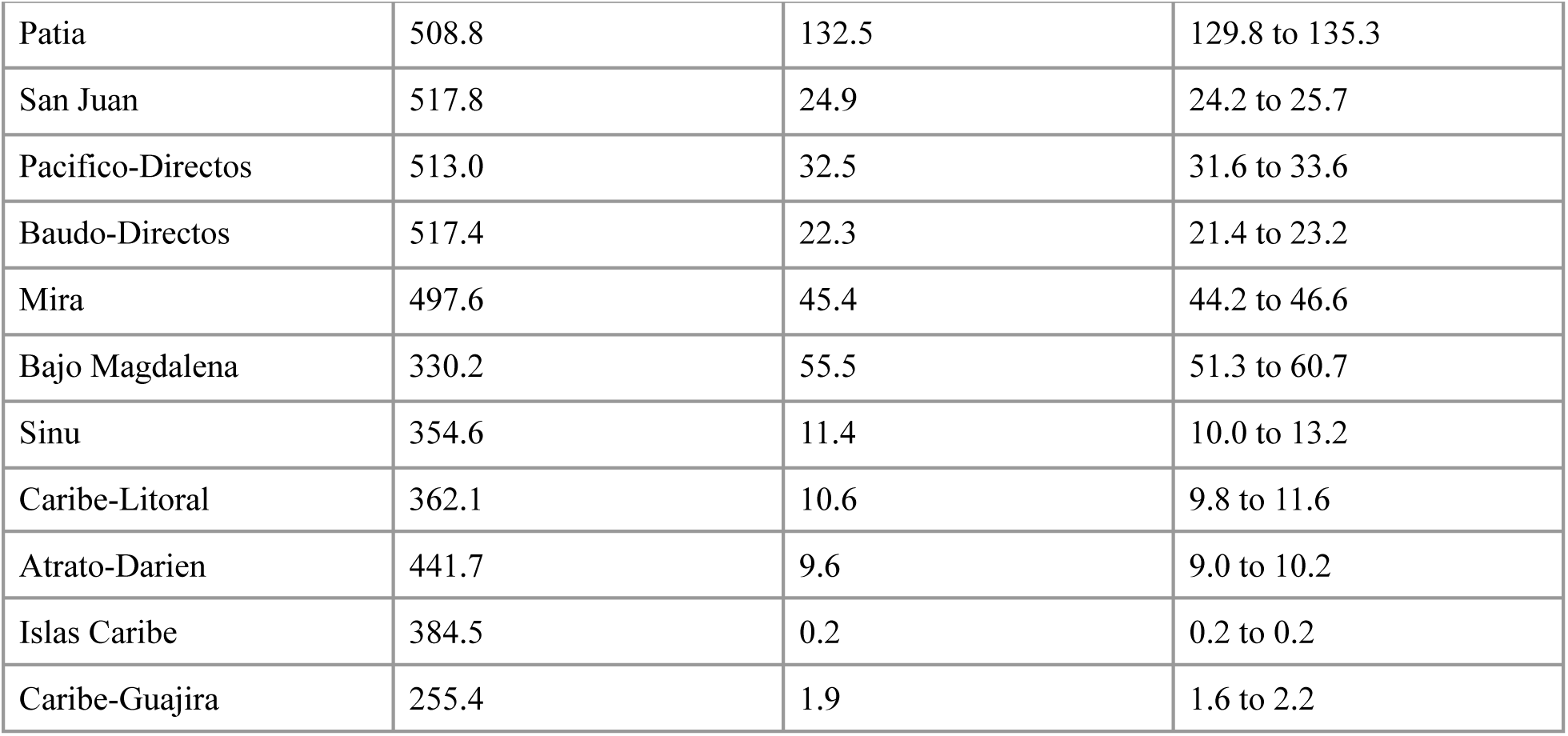
Zone-level Monte Carlo carbon stock results.

| Zone | Carbon Density (Mg C/ha) | Total CO<sub>2</sub>e (millions) | 95% CI (millions) |
| --- | --- | --- | --- |
| Tapaje-Dagua-Directos | 528.1 | 131.6 | 129.0 to 134.4 |
| Patia | 508.8 | 132.5 | 129.8 to 135.3 |
| San Juan | 517.8 | 24.9 | 24.2 to 25.7 |
| Pacifico-Directos | 513.0 | 32.5 | 31.6 to 33.6 |
| Baudo-Directos | 517.4 | 22.3 | 21.4 to 23.2 |
| Mira | 497.6 | 45.4 | 44.2 to 46.6 |
| Bajo Magdalena | 330.2 | 55.5 | 51.3 to 60.7 |
| Sinu | 354.6 | 11.4 | 10.0 to 13.2 |
| Caribe-Litoral | 362.1 | 10.6 | 9.8 to 11.6 |
| Atrato-Darien | 441.7 | 9.6 | 9.0 to 10.2 |
| Islas Caribe | 384.5 | 0.2 | 0.2 to 0.2 |
| Caribe-Guajira | 255.4 | 1.9 | 1.6 to 2.2 |

The ranking reflects a tension between ecological potential and operational accessibility: the highest-scoring zones (Patia, Tapaje-Dagua) are, arguably, among the least accessible due to security constraints and remoteness, while lower-scoring Caribbean zones offer easier access.

### 4.5 REDD+ White Space Assessment

Of 276,430 ha nationally, 133,794 ha (48.4%) qualify as white space with no existing REDD+ project claims (Figure 9). The remaining areas were classified as: PROTECTED_MEDIUM_ADDITIONALITY (732 polygons, 126,504 ha), CONTESTED (11 polygons, 3,132 ha, in the Sinu zone overlapping Vida Manglar), or ADJACENT_LEAKAGE (1 polygon).

**Figure 9.**
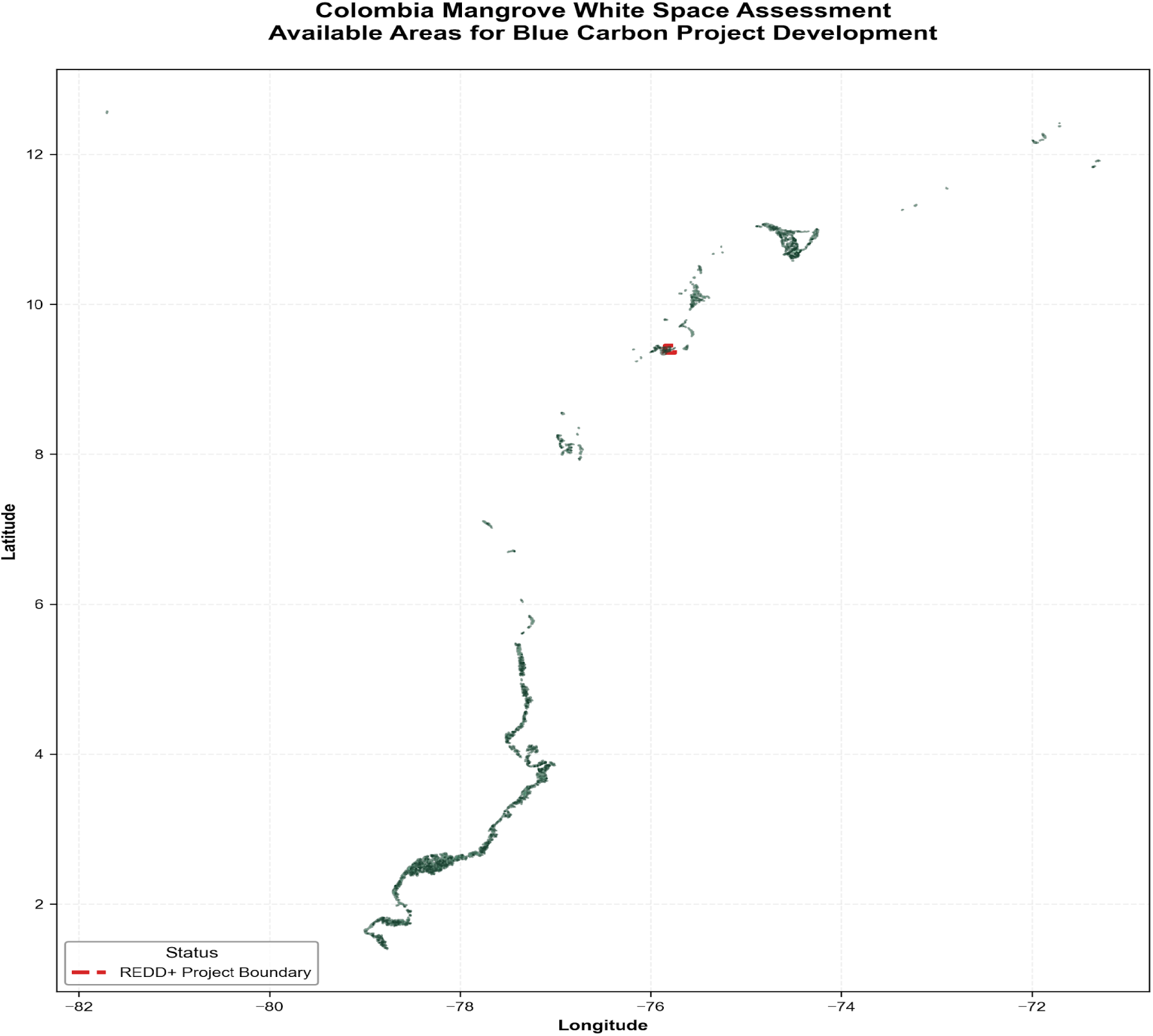
REDD+ White Space Distribution. Green: white space (133,794 ha, 48.4%). Purple/sage: protected areas with medium additionality (126,504 ha, 45.7%). Red: contested areas (3,132 ha). White space concentrated on the Pacific coast (114,000 ha, 85% of total). Source: Verra VCS registry (February 2026), RUNAP, IDEAM. CRS: EPSG:4686.

White space is predominantly located on the Pacific coast, accounting for 114,000 hectares (85% of the total). A critical distinction is that the hydrographic zone boundaries utilized in the REDD+ assessment are more extensive than the classified mangrove area employed for carbon stock calculations. For instance, the San Juan white space area (35,724 ha) is defined by the hydrographic zone limit, whereas the corresponding carbon analysis is restricted to the 13,102 hectares of classified mangrove polygons situated within that zone.

The assessment identified 11 polygons, encompassing 3,132 ha, within the Sinu zone as contested. This finding aligns with the area reported by Conservation International’s Vida Manglar project (VCS 2290) as potentially overlapping. Notably, only 689 ha (7.0%) of the Sinu zone qualifies as “white space.” The observed discrepancy between the 3,132 ha overlap and the project’s reported 7,561 ha total area is attributable to two factors: the use of an approximate bounding box geometry in the assessment rather than the project’s precise KML boundary, and the inclusion of non-mangrove habitats within the overall project area.

### 4.6 ML Emulator Performance

The CatBoost emulator achieves R² = 0.926 (RMSE = 21.3 Mg C/ha) on GroupKFold cross-validation (Figure 10), outperforming XGBoost (R² = 0.758), Random Forest (R² = 0.629), and Ridge regression (R² = −0.223) (Figure 11).

**Figure 10.**
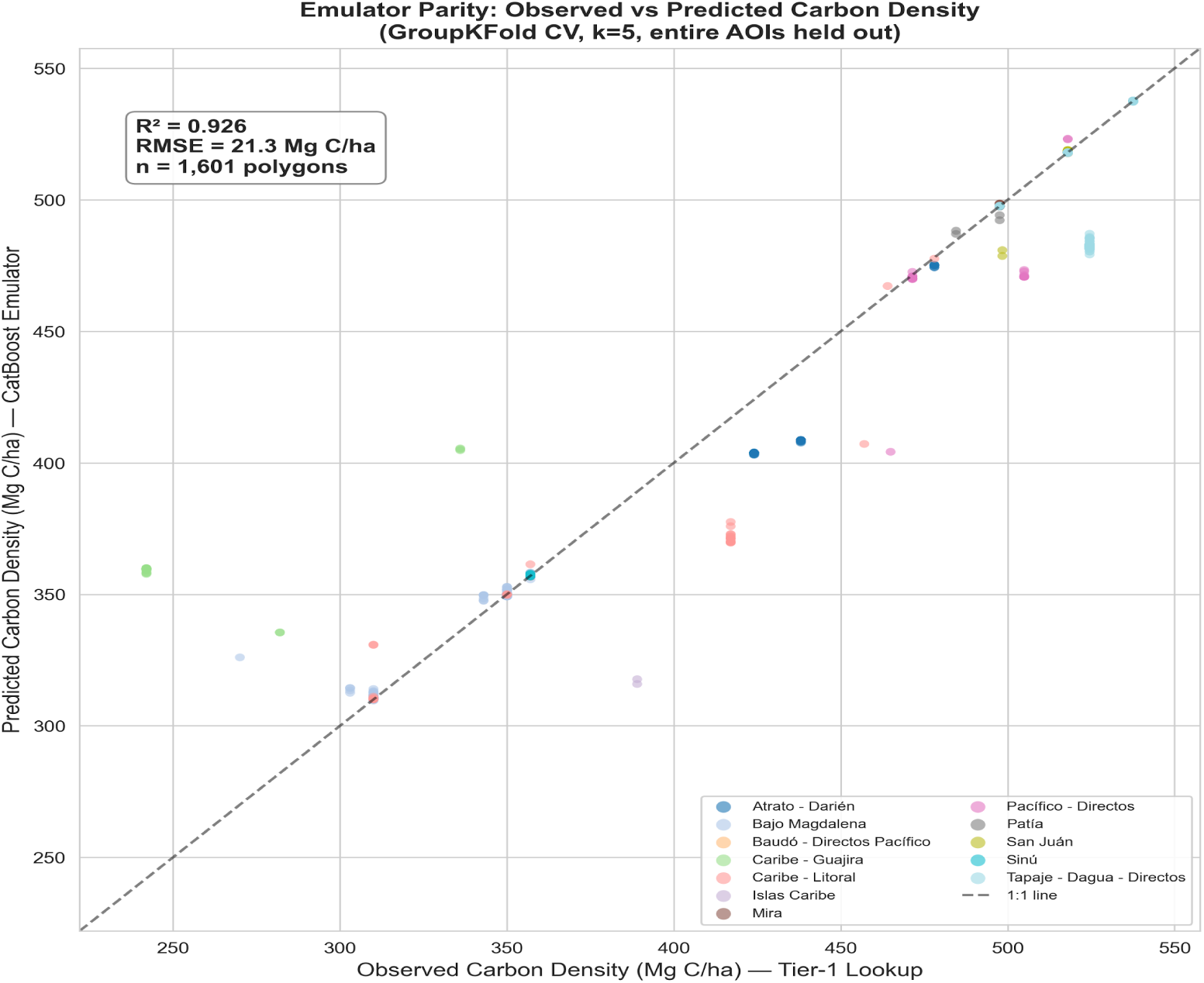
CatBoost Emulator Parity Plot. Scatter plot of emulator-predicted vs. Tier 1 lookup-derived carbon density for 1,601 polygons. GroupKFold CV (k = 5): R² = 0.926, RMSE = 21.3 Mg C/ha. Dashed: 1:1 line. Points colored by the hydrographic zone.

**Figure 11.**
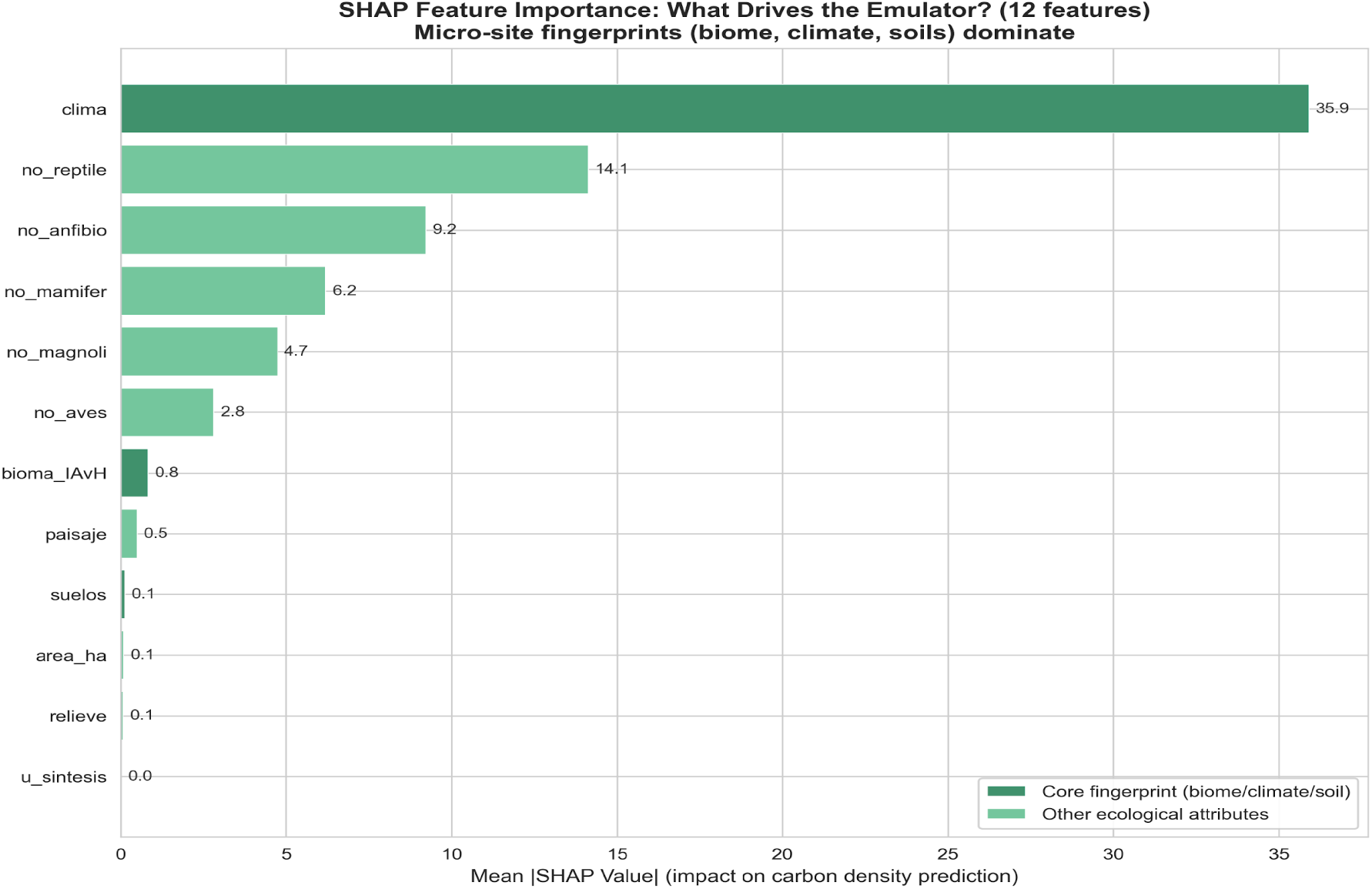
SHAP Feature Importance Fingerprint. Horizontal bar chart of mean absolute SHAP values for all 12 features. Climate class (clima) dominates (35.9, 47.4%). Biodiversity counts contribute 48.9%. Top 3 features explain 78.2%; top 5 explain 92.6%.

The SHAP analysis (Figure 11) establishes a robust hierarchy of feature importance influencing carbon density variation. Climate classification emerges as the preeminent predictor, singularly accounting for 47.4% of the observed variance. The aggregation of biodiversity counts across five distinct taxonomic groups contributes an additional 48.9%, bringing the cumulative explanatory power of climate and biodiversity to 96.3%. In contrast, the biome classification utilized for the Tier 1 AGB lookup contributes a marginal 0.8%. This significant disparity demonstrates that the granular stratification provided by climate class and biodiversity metrics offers superior predictive resolution compared to the broader biome category. Critically, this result provides empirical validation for the climate-stratified SOC approach, as the emulator independently identifies climate as the dominant variable, confirming that the six-class climate stratification effectively captures the principal axis of carbon density variation.

Quantile regression (P10/P90) demonstrates that the per-polygon uncertainty exhibits a distinct spatial structure (Figure 12). Specifically, polygons located within climatically homogeneous Pacific zones display narrow uncertainty bands. In contrast, polygons situated at zone boundaries or within ecologically transitional areas show wider bands, effectively identifying these as priority targets for subsequent field verification efforts.

**Figure 12.**
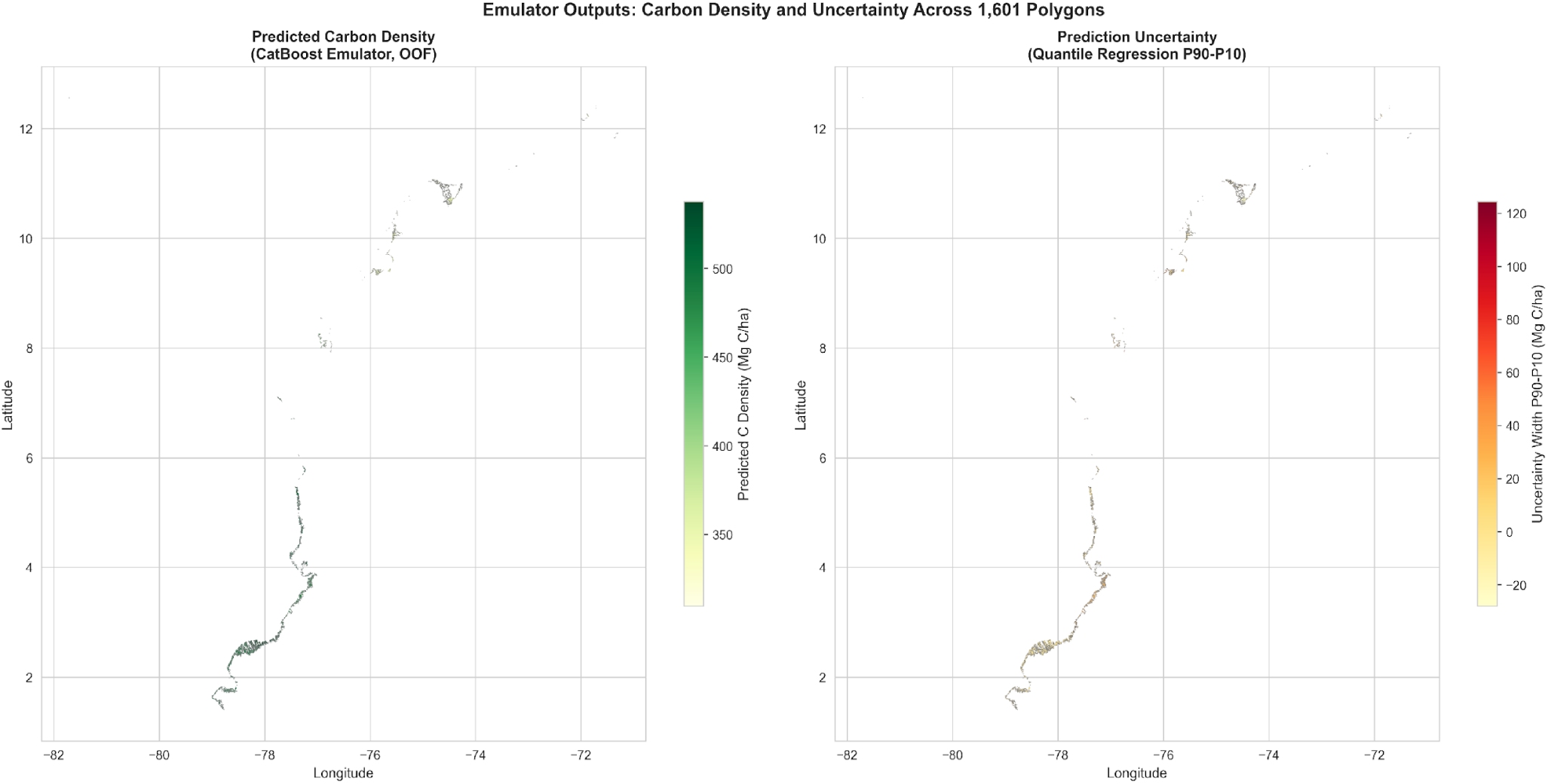
Spatial Uncertainty Quantification. Per-polygon uncertainty (P90 minus P10) from CatBoost quantile regression. Wider bands (warm colors): high uncertainty at zone boundaries. Narrow bands (cool colors): well-constrained Pacific zones. CRS: EPSG:4686.

### 4.7 Constraint-Filtered Portfolio Analysis

The initial theoretical assessment of blue carbon opportunity identified a substantial white space of 133,794 hectares (ha). However, the rigorous application of critical constraints, specifically governance, security, and regulatory factors, necessitated a significant filtering process, resulting in a substantially reduced and operationally viable portfolio of 4,000 to 12,000 ha. This refined range constitutes only 3% to 9% of the initial theoretical opportunity, underscoring the critical dependence of operational feasibility on stringent site intelligence. Security considerations emerged as a primary constraint, eliminating several zones that presented high theoretical potential. For instance, the Patia zone, despite possessing the highest theoretical white space (28,466 ha) and optimal TOPSIS score, is situated within an environment characterized by active armed conflict, illicit coca production, and established narco-trafficking corridors. Similarly, the Atrato-Darien area has suffered a correlation between mangrove extent loss (30% over six years) and the operational control exerted by armed non-state actors. The San Juan zone is further compromised by the inclusion of Buenaventura, a well-documented international hub for narco-trafficking activities.

Regulatory and land-use overlaps also significantly constrained the viable portfolio. The Sinu zone is heavily dominated by the existing *Vida Manglar* project (VCS 2290), leaving a residual white space of only 689 ha, representing a minimal 7.0% of its total theoretical opportunity. In Bajo Magdalena, the significant white space (15,892 ha) extensively overlaps with the Cienaga Grande de Santa Marta protected area complex. The primary, binding constraints here are the undefined legal status of carbon rights within this protected area and the critical requirement for extensive, non-negotiable hydrological restoration before project development. Consequently, viable, near-term site selection is strictly governed by pre-defined governance readiness criteria. Successful sites must demonstrate the presence of established, formalized community governance structures (specifically, *Consejos Comunitarios* with legally recognized land tenure), provide tangible evidence of pre-existing and stable conservation partnerships, and be verified as operating under stable security conditions. The application of these essential governance and security filters is the decisive factor that reduced the initial potential to the final, pragmatic portfolio of 4,000 to 12,000 ha.

### 4.8 Representative Site Typologies

The utility of the HiGEBCA pipeline is demonstrated through the systematic characterization of four distinct mangrove typologies, each providing empirical evidence regarding unique contextual challenges and opportunities critical for the successful implementation of blue carbon initiatives.

1. Caribbean Urban Mangrove (Caribe-Litoral zone, Cartagena): This location serves as a methodology demonstration site, covering 1,524 hectares. It is defined by high logistical accessibility, established community governance (Consejo Comunitario), and documented, active restoration efforts, as evidenced by the planting of over 11,000 trees.
2. Pacific Estuarine Typology (Tapaje-Dagua-Directos): This typology has been quantitatively identified as the highest-carbon-density opportunity, registering a TOPSIS score of 0.634 and a density of 528.1 Mg C/ha. It encompasses significant white space (50,180 ha) that is currently supported by established community-based conservation programs.
3. Caribbean Deltaic Typology (Bajo Magdalena): This case exemplifies a scenario where fundamental anthropogenic infrastructure, specifically, the severance of hydrological connectivity since 1956, acts as a primary constraint on restoration viability, regardless of the zone’s inherent ecological potential.
4. High-Potential Exclusion Typology (Patia): Characterized by a high ecological score (TOPSIS 0.672) and significant white space (28,466 ha), this zone serves to quantify the critical disparity between theoretical ecological potential and operational viability. It empirically demonstrates how active armed conflict renders the area inaccessible, thereby precluding project implementation.

## 5. Discussion

### 5.1 Comparison to Existing Blue Carbon Assessments

Standard blue carbon site assessments typically rely on five or fewer ecological attributes per spatial unit, often applying coast-level or regional carbon factors to satellite-derived extent estimates. For example, the most comparable published assessment employed spatial Multi-Criteria Decision Analysis (MCDA) with only three criteria (Rog et al. 2024). In contrast, the HiGEBCA pipeline developed herein incorporates 47 attributes per polygon, alongside biome-specific parameters for above-ground biomass (14 lookup values), climate-stratified soil carbon (6 classes), and integrated biodiversity co-benefit quantification derived from government-sourced field data across five taxonomic groups. A recent synthesis documented that methodological differences in blue carbon science can lead to carbon stock estimates differing by up to 10-fold across studies (Macreadie et al. 2025). The HiGEBCA polygon-level approach, which propagates uncertainty through Monte Carlo simulation and matches carbon estimation parameters to the specific ecological context of each polygon rather than relying on regional averages, significantly reduces this variance.

### 5.2 Governance Constraint Integration as Methodological Innovation

Critically, no published blue carbon assessment reviewed has systematically integrated governance feasibility into site prioritization; standard approaches typically treat governance as a post-hoc qualifier after ecological potential is assessed. In contrast, the HiGEBCA pipeline rigorously incorporates governance, security, and regulatory constraints as analytical layers, applying the same methodological rigor used for ecological characterization. This integration yields a constraint-filtered portfolio analysis that explicitly quantifies opportunity loss at each filtering stage: the initial national estate of 276,430 ha is sequentially reduced to 133,794 ha (REDD+ white space) and finally to the operationally accessible range of 4,000 to 12,000 ha. This transparency is crucial, preventing the overstatement of accessible opportunity, which is a common failure mode in blue carbon project development.

### 5.3 Scalability and Replicability

The HiGEBCA pipeline is explicitly designed for cross-country replication, with data requirements comprising: (1) a national or regional ecosystem map detailing polygon-level ecological attributes, (2) a geospatial registry of REDD+ projects, and (3) a protected area registry including boundary data. Several Latin American countries possess equivalent foundational mapping infrastructure, exemplified by Ecuador’s national mangrove census (161,835 ha), Panama’s UK-funded Blue Natural Heritage initiative, and Mexico’s CONABIO ecosystem mapping. Ecuador serves as a particularly strong scalability test case. The country maintains 161,835 ha of mangroves, with 42.85% formally managed under community custodianship via *Acuerdos de Uso Sustentable y Custodia del Manglar* (AUSCM) agreements. The national census has documented 26 consecutive years of mangrove area stability, which constitutes some of the strongest empirical evidence in the Americas demonstrating that community-governed coastal ecosystems can achieve management outcomes equivalent to, or superior to, those of state management. The regulatory pathway is established by Ecuador’s memorandum of understanding with Gold Standard for a national blue carbon methodology, signed on 17 July 2025.

### 5.4 Methane Considerations

While blue carbon ecosystems like mangroves are globally recognized for their substantial capacity to sequester atmospheric carbon dioxide, a critical and often-overlooked factor in accurate carbon accounting is the emission of methane (CH_4_). Methane is a potent greenhouse gas, and its release from these environments, particularly through plant stems, can potentially offset a significant portion of the carbon sequestration benefit. Therefore, understanding and accurately quantifying the balance between carbon burial and CH_4_ emissions is crucial for establishing credible blue carbon digital Measurement, Reporting, and Verification (dMRV) protocols and ensuring the climate benefits of these ecosystems are not overstated. Recent research documented that mangrove tree stem methane emissions offset 16.9% of sediment carbon burial at the global scale (Qin et al. 2025). However, this finding has been contested; a reanalysis removing statistical outliers produced a revised offset estimate of 0.5% to 4.7% when soil methane oxidation processes are appropriately accounted for. Consequently, conservative carbon accounting necessitates a methane prudence buffer of 15% to 25% until methodology evolves. Molecular approaches, specifically shotgun metagenomics, present a promising path toward accurate, site-specific methane accounting. Untargeted metagenomic sequencing allows for the direct profiling of microbial community functional gene content, enabling the quantification of the ratio of methanogenic (*mcrA*) to methanotrophic (*pmoA*) genetic capacity. The Microflora Danica atlas, an analysis of 10,683 metagenomes (Singleton et al. 2025), has been instrumental in identifying numerous previously uncharacterized methanotrophic species that appear to dominate methane oxidation in undisturbed ecosystems. These novel species are absent from current reference databases and are thus undetectable by conventional amplicon-based surveys. Crucially, only shotgun metagenomics can detect new functional genes without reliance on prior reference sequences, establishing it as a candidate technology for next-generation blue carbon dMRV.

### 5.5 Implications for Blue Carbon MRV and Market Context

The HiGEBCA pipeline establishes a robust, multi-layered architectural framework for a comprehensive, full-stack digital Measurement, Reporting, and Verification (dMRV) system. This system integrates a foundational geospatial layer, which precisely delineates carbon stock distribution and governance, with advanced monitoring and verification components. Specifically, the planned integration of satellite-derived change detection, utilizing Normalized Difference Vegetation Index (NDVI), Normalized Burn Ratio (NBR), and Ratio Vegetation Index (RVI) from Sentinel-2, will transition the static geospatial characterization into temporal monitoring capabilities at a 10-meter resolution. Furthermore, the inclusion of molecular verification methods, specifically, environmental DNA (eDNA) metabarcoding for taxonomic inventory and shotgun metagenomics for functional profiling, will provide crucial empirical evidence regarding microbially-mediated sediment biogeochemistry at designated sites.

The broader market context is defined by structural supply constraints, as evidenced by the S&P Global Platts DBC-1 blue carbon benchmark reaching $29.30/tCO₂e in August 2025. Projects that achieve verified co-benefits demonstrably command price premiums. For instance, Sylvera (2026) reports that credits from projects scoring 5 on co-benefit metrics average $25 per credit, with certification under CCB Standards or SD VISta yielding premiums exceeding 50% relative to unverified projects. Furthermore, the estimated national carbon stock of 478 million tCO₂e represents a substantial theoretical asset value, ranging from $9.6 to $21.5 billion based on a valuation range of $20 to $45 per tCO₂e; however, the actual accessible stock, after accounting for practical constraints, is likely to be a fraction of this total. Consequently, the comprehensive ecological characterization provided by the HiGEBCA pipeline, as presented herein, is directly relevant for achieving the highest tiers of co-benefit certification, thereby maximizing both the project’s conservation outcomes and its community impact.

The HiGEBCA pipeline offers a robust solution for enhancing the credibility of nature-related financial disclosures and addressing structural critiques within carbon markets, such as those related to REDD+. Specifically, its outputs, including biodiversity baselines, carbon estimates with quantified uncertainty, and analyses of governance constraints, can provide invaluable inputs for double-materiality reporting under the European Union Corporate Sustainability Reporting Directive (CSRD) and European Sustainability Reporting Standards (ESRS) E4. Critically, the pipeline directly confronts the “disconnected paper realities” problem prevalent in carbon project documentation, where ecological conditions are often described through generic, unmeasurable narratives. The HiGEBCA pipeline counters this by grounding its carbon estimates in a 47-attribute-per-polygon characterization, which leverages government-verified ecological classifications (e.g., biome, soil, climate, biodiversity). This approach replaces reliance on project-developer-generated narratives with scientifically verifiable data. Numerical evidence from the CatBoost emulator further validates this rigor, demonstrating that biodiversity counts explain 48.9% of the variation in carbon density, thereby establishing that ecological reality and carbon storage are intrinsically coupled dimensions, not independent constructs assembled merely for reporting compliance.

A significant challenge in traditional carbon accounting is the “static baseline” problem, where baselines are treated as fixed at the project’s start, despite the continuous evolution of ecological conditions. To address this, the HiGEBCA pipeline incorporates an emulator architecture that supports annual baseline updating. Specifically, as new ecosystem map editions become available (typically every 2–4 years via releases from IDEAM), the Monte Carlo carbon estimation and CatBoost model can be retrained to accurately reflect current conditions. This dynamic approach directly addresses the methodological gap identified in next-generation REDD+ protocols, which advocate for dynamical baselines on 2–3 year reassessment cycles. Furthermore, the planned integration of satellite-derived change detection data (utilizing NDVI, NBR, and RVI from Sentinel-2 at 10 m resolution) will facilitate sub-annual monitoring of canopy condition. This capability can move mangrove dMRV closer to the temporal resolution of established forest REDD+ protocols, while simultaneously operating at a superior spatial resolution.

Addressing the pervasive problem of transparency, where current carbon project documentation often comprises hundreds of pages of PDF reports that hinder, rather than facilitate, accurate methodological assessment and informed decision-making, the HiGEBCA pipeline introduces a verifiable and reproducible infrastructure. This system ensures full code availability and complete data provenance. Consequently, every output, from the carbon estimates and TOPSIS scores to the selection of field sampling candidates, is fully traceable to its original government-source input data via explicitly documented transformations. This commitment to data lineage and transparency is designed to align with the stringent demands of emerging blockchain-based carbon registries, which require cryptographic verification of data history, although the current implementation does not incorporate blockchain technology.

While these methodological features do not resolve all issues inherent to carbon market credibility, such as demonstrating additionality, quantifying leakage, and ensuring long-term permanence, they fundamentally establish a transparent, ecologically-grounded, and dynamically-updated framework for baseline assessment, directly addressing known shortcomings in current blue carbon practice.

### 5.6 Limitations

Several limitations constrain the current analysis and warrant careful interpretation of the results. First, the carbon estimates rely exclusively on Tier 1 methodologies. All carbon stock values are literature-based (Tier 1 per IPCC 2013 Wetlands Supplement definitions), derived from published allometric relationships and regional Soil Organic Carbon (SOC) values rather than site-specific field measurements. Although the Monte Carlo uncertainty propagation quantifies parametric uncertainty, it does not account for systematic bias inherent in Tier 1 assumptions. Consequently, field validation is a prerequisite before these estimates can support carbon credit issuance, a necessity addressed by the optimized field sampling design detailed in Section 3.8. Second, the biodiversity data is derived from government sources. Species richness counts originate from the national ecosystem map’s coded attributes, which reflect established government mapping protocols rather than independent field surveys or direct species observations. The codes represent assemblage types associated with each polygon’s ecological classification. While beta diversity patterns (i.e., compositional differences between zones) remain robust, the absolute species counts should be interpreted as relative indicators of potential richness, rather than census data. Third, the analysis presents a static temporal snapshot based on the 2024 map edition, while mangrove ecosystems are inherently dynamic, influenced by factors such as Pacific El Niño-Southern Oscillation (ENSO) cycles affecting productivity and extent, ongoing deforestation in conflict zones, and changes in canopy coverage due to restoration programs. A planned extension will incorporate satellite-based change detection to address this temporal constraint. Fourth, the TOPSIS prioritization relies on expert-assigned weights, representing one defensible configuration. Alternative weight configurations would generate different rankings. However, the top-ranked zones (Patia, Tapaje-Dagua) demonstrate robustness to moderate weight perturbations due to their consistently high scores across multiple criteria, whereas mid-ranked zones exhibit sensitivity to the relative weighting of area versus ecological condition metrics. Fifth, the source map resolution introduces spatial uncertainty. The 1:100,000 scale corresponds to a positional accuracy of approximately 100 meters. Therefore, fine-grained spatial analyses, such as patch boundary delineation or edge effects below 50 meters, must be interpreted cautiously, acknowledging this resolution constraint. Sixth, the white space assessment for conservation opportunities identified only one registered mangrove REDD+ project (Vida Manglar). The potential existence of additional projects under different registries or in earlier development stages suggests that the calculated 133,794 ha white space figure may represent an overestimate of the currently available, unregistered area. Finally, the landscape metrics terminology warrants clarification. The HiGEBCA pipeline computes landscape metrics analogous to those defined by FRAGSTATS (McGarigal et al. 2012); however, the metrics (core area integrity, fragmentation risk, isolation index) are custom implementations computed in Python using the GeoPandas library, not the FRAGSTATS software directly.

## 6. Conclusion

The HiGEBCA pipeline is explicitly designed as a reproducible infrastructure. Its data requirements, specifically polygon-level ecological mapping, a REDD+ project registry, and protected area boundaries, are anticipated to be readily available in numerous Latin American countries. The entire methodology is thoroughly documented, fully reproducible from its source data, and intentionally extensible to incorporate future layers such as satellite-based change detection and molecular verification data. However, to advance high-integrity blue carbon assessments, future work should prioritize: (1) field validation of Tier 1 carbon estimates within the 50 priority polygons identified by the optimal sampling design; (2) integration of satellite-derived change detection for continuous temporal monitoring; (3) cross-country replication, commencing with Ecuador; and (4) the integration of molecular verification techniques (e.g., eDNA biodiversity inventories and metagenomic functional profiling) to enhance site-specific Measurement, Reporting, and Verification (MRV) protocols.

## Acknowledgments

The primary dataset used in this analysis is maintained by IDEAM, SINCHI, IAvH, and INVEMAR as part of the Colombian National Ecosystem Map program. The authors gratefully acknowledge these institutions for producing and maintaining the 1:100,000-scale ecological mapping infrastructure that makes polygon-level analysis possible. Protected area boundary data were obtained from the Colombian RUNAP database. REDD+ project boundary data were obtained from the Verra VCS public registry.The HiGEBCA pipeline was developed using an agentic workflow architecture in which the author orchestrated multiple large language model agents as specialized research assistants across the full analytical lifecycle. AI agents were directed to generate and iteratively refine geospatial processing scripts, Monte Carlo simulation code, machine learning pipelines, and visualizations. All analytical decisions, parameter selections, data interpretations, and scientific conclusions are the sole responsibility of the author.

## Data Availability

The primary dataset (IDEAM 1:100,000-scale National Ecosystem Map, 2024 edition) is maintained by the Colombian government and available through IDEAM’s public data portal. Processed analytical outputs (CSV files, GeoJSON geometries, interactive visualizations) are available from the corresponding author upon request. REDD+ project boundaries are publicly available from the Verra VCS registry. Protected area boundaries are publicly available from Colombia’s RUNAP database.

## Code Availability

The full analytical pipeline (15 modules across 4 Jupyter notebooks, Python scripts for notebook generation, and a FastMCP server for interactive screening) is available from the corresponding author upon request. The pipeline uses open-source libraries: GeoPandas, CatBoost, SHAP, scikit-learn, Matplotlib, and Plotly.

